# Homing pigeon (*Columba livia*) dominance hierarchies are stable over time and resistant to perturbation

**DOI:** 10.1101/2022.04.18.488671

**Authors:** Amaia A. Urquia-Samele, Steven J. Portugal

## Abstract

Dominance hierarchies are a social dynamic common in many species, which help balance the costs and benefits of social living. Which individuals in a group occupy high ranking positions is influenced by a multitude of different intrinsic and/or extrinsic factors. While homing pigeons (*Columba livia*) have been a model system for their navigational abilities and flight dynamics, less is known about their dominance hierarchies and general social behaviour when not in flight. Here we document the dominance hierarchies in two closed populations of homing pigeons, and investigate the physiological factors associated with dominance rank, including body size, metabolic rate, personality, and iridescent plumage % reflectance. Measurements of body size and resting metabolic rate all positively correlated with dominance rank in accordance with the *performance model* of energetic management. Iridescent plumage % reflectance was negatively correlated with rank, suggesting a potential cost of dominance. Attempts to alter hierarchy structure through manipulations of body mass and feather iridescence were ineffective which hierarchy dynamics remaining stable throughout the perturbations and across measurement sessions.

## INTRODUCTION

Living as part of a group is a common occurrence in many animal species (Elgar 1989). Being part of a collective group can come with many benefits, such as increased protection from predation (Bertram 1978), opportunities for social learning (Slagsvold and Wiebe 2011), higher foraging success (Alexander 1974), energetic savings (Markham and Gesquiere 2017) and increased mating opportunities (Ulrich *et al*. 2018, Elgar 1989). Conferring these benefits can, however, still come at a cost. Such costs can include increased transmission of diseases (Witte *et al*. 2020), food being stolen by conspecifics (Brockmann and Barnard, 1979), and an unequal sharing of resources (Alexander 1974). Due to competition, and this unequal sharing of resources, conflicts can arise in animal groupings. Heightened competition can be costly, as it typically involves high energy expenditure and/or the risk of injury, with potentially no pay off from the aggressive interaction if unsuccessful (Alexander 1974). In order to reduce these potential costs, individuals may avoid conflict with conspecifics they have previously lost fights with to avoid wasting energy, and this frequently leads to the formation of dominance hierarchies (Hand 1986, Drews 1993).

Dominance is a measure of status within a group and is typically – but not always – determined based on aggressive interactions. Aggressive individuals are able to win more interactions and typically procure a higher proportion of resources, and a higher dominance rank. Dominance hierarchies can have differing structures, ranging from differing degrees of linearity to triangular or despotic arrangements, depending on the species (de Vries, Stevens and Vervaecke 2006). When dominance hierarchies are highly linear, they can be correlated with certain physical or intrinsic traits (Beacham 2003). Common parameters typically associated with determining dominance ranks include high body mass and/or structural size, which greatly aids defence of resources and can also assist in resistance to environmental stresses (Haramis *et al*. 1986, Moreno-Opo, Trujillan and Margalida 2020). This positive relationship between body mass and rank is documented in a diverse range of species such as in zebra finches *Taeniopygia guttata* (Cuthill *et al*. 1997), masu salmon *Oncorhynchus masou* (Yamamoto, Ueda and Higashi 1998) tuataras *Sphenodon punctatus* (Moore *et al*. 2007) and white-tailed deer *Odocoileus virginianus* (Taillon and Côté 2006). Other variable factors which are reported to relate to dominance rank include personality traits such as boldness and neophobia, (Fox *et al*. 2009) and metabolic rate (Bryan and Newton 1994). Unlike the relationship between rank and body mass, the association between personality, metabolism and rank is typically more complex. Several energetic-based models predict the potential relationships between metabolic rate and personality traits, and in turn dominance, under various scenarios (Mathot, Dingemanse and Nakagawa 2019). The *allocation model* suggests variation in metabolic rate is not associated with changes in behaviours associated with energy gain; individuals with higher metabolic costs are less likely to invest in foraging behaviours and overall boldness personality traits, due to their energetically costly nature (Mathot, Dingemanse and Nakagawa 2019). This relationship between metabolic rate and behaviour has been observed in species such as mosquito fish *Gambusia affinis* (Cote *et al*. 2010), deer mice *Peromyscus maniculatus* (Careau *et al*. 2011), Ouachita dusky salamanders *Desmognathus brimleyorum* (Gifford, Clay and Careau 2014) and zebra finches *Taeniopygia guttata* (Mathot *et al*. 2009). In contrast the *performance model* predicts that metabolic rate correlates with increased of behaviours involved in energy gain. Under this model, individuals will have greater activity if their metabolic rate is higher, due to their energetic requirements (Mathot, Dingemanse and Nakagawa 2019).

This interaction between behaviour and metabolism can be highly influential in the determination of dominance hierarchies, with certain personality types being more common in certain rank positions. In several species such as common lizards *Zootoca vivpara* (Le Galliard *et al*. 2013), pied flycatchers *Ficedula hypoleuca* (Røskaft *et al*. 1986), great tits *Parus major* (Røskaft, *et al*. 1986) and mountain chickadees *Poecile gambeli*, for example, exploratory birds will occupy higher ranking positions, while opting to kleptoparasite subordinates instead of investing in foraging behaviours (Fox *et al*. 2009, Brown, Jablonski and McCormack 2007).

Regardless of what factors are determining a dominance hierarchy, once a hierarchy has been established, aggression and ranks can remain relatively stable, with aggression levels remaining lower than before the hierarchy was formed (Hewitt, Macdonald and Dugdale 2009, Portugal *et al*. 2020). However, this stability can change over time due to factors such as seasonal modifications in physiology and behaviour (Forand, Marchinton and Miller 1995), immigration and emigration, or the death of a high ranking member (Grossel *et al*. 2021). As many dominance hierarchy types can be dependent on the body condition of the individuals, any changes in body condition of a dominant individual, such as body mass, may lead to disruption of the hierarchy and group dynamics (Robinson 1986). For example, artificial mass loading of subordinate homing pigeons *Columba livia* saw a disruption of hierarchy stability, with lower ranking individuals gaining rank after the increase in body mass (Portugal *et al*. 2020). This was despite the hierarchy being stable for a number of years prior to the artificial mass manipulation experiments.

Homing pigeons are a model system for the study of dominance and general social dynamics (Bouchard, Goodyer and Lefebvre 2006), navigational ability (Taylor, Portugal and Biro 2017) and intelligence (Carter and Werner 1978). Homing pigeons are a domestic form of rock doves which have been selectively bred for their flight speed and navigation over centuries (Matthews 1951, Shapiro and Domyan 2013). Previous investigations into the dominance hierarchies in pigeon social groups have indicated a strong linear dominance hierarchy, strongly correlated with body mass (Portugal *et al*. 2020). It has also been suggested that elements of plumage iridescent play an important role in flock social structure and breeding display behaviour (Johnston 1992). Using two flocks of captive homing pigeons, we tested a series of hypotheses pertaining to the underlying traits, such as body mass, metabolic rate and personality, which may determine position within a dominance rank, before perturbating certain traits to determine how stable these rank positions are. Specifically, we first tested whether body mass, structural size, personality, resting metabolic rate or the degree of plumage iridescent predicted dominance rank. Thereafter, we manipulated aspects of body mass and plumage reflectance, to determine whether hierarchy stability was susceptible to perturbation when these traits where artificially manipulated.

## METHODS

### Birds

Homing pigeons (*n =*33) were housed at Royal Holloway University London (Egham, UK), split between two lofts (dimensions 3.6m x 2.4m). Birds had unrestricted access to food (Johnstone &Jeff Four Season Pigeon Corn, Gilberdyke, UK), grit and water. Loft 1 comprised 17 individuals (8 males, 9 females) and loft 2 sixteen (8 males and 8 females). Sex was determined by genetic testing of feather samples (Animal Genetics, UK). While no birds were added to either loft during the duration of the experiment, two pigeons were omitted part way through the study period from loft 2 due to (unrelated) injuries sustained after the first dominance control study. In total, 1 male and 1 female were removed. Individual pigeons were weighed using scales (PS1C EPOS integrated scales, Swansea) to the closest 10g. Structural size was determined through measuring their tarsus length (from the elbow joint to the ankle joint) using electronic callipers (Louisware from Amazon).

### Dominance hierarchies

The dominance hierarchies of each of the two pigeon flocks was determined through observing aggressive interactions, following protocols of Portugal *et al*. (2017a) and Portugal *et al*. (2017b). Food was removed from the lofts the day before each dominance trial, at approximately 15:00. Water remained available at all times. Before each dominance trial, all birds were caught and placed into a pigeon carrier. Each bird was then labelled with back mounted Velcro (VELCRO®) fixed in place with epoxy resin (Araldite® two part epoxy). A single small feeder, half filled with food, was placed against the wall of the home loft opposite the carrier. The placement of the feeder against the wall removed several access points for the birds, encouraging confrontation. Ceiling mounted cameras (GoPro Hero4 or Akaso EK7000) recorded interactions across the loft over a 60-minute period, once the pigeons were released from the carrier. Upon completion of each dominance trial, food was returned to loft.

The initial dominance hierarchies trials (*n =* 10) of the two flocks were studied over a 2-week period across February and March 2021. A second control of 10 dominance trials was conducted in May and June 2021. These trials were undertaken to determine if any carry over effects from the first manipulation trials were evident. During dominance trials, all aggressive interactions were recorded, included pecking, chasing, wing slapping, neck grabbing and beak grabbing (see Portugal *et al*. 2020 for full definitions). Which pigeon was the aggressor (winner) and which pigeon was the subject (loser) were recorded using BORIS free software (Friard and Gamba 2016). Dominance information was then displayed in a matrix for each dominance trial and used to calculate dominance rank using David’s score (see Gammel *et al*. 2003).

### Personality assays

Individual personality was quantified by establishing individual tendency for lone exploration in an unfamiliar environment (Portugal *et al*. 2017a). Exploration trials took place in February and March 2021. For exploration, individual birds were taken from their home loft and placed in an animal carrier. The carrier was placed in the corner of an uninhabited loft which had been divided into 3 equal sections with tape to mark three respective zones (see Portugal *et al*. 2017a). The pigeon was released from the carrier and allowed to explore the loft for 30 minutes. The duration spent in the carrier, zone one, zone two, zone three, as well as perches, was recorded. Five repeats were carried out per pigeon, with each repeat for each pigeon taking place approximately every 2 days; no repeats on the same pigeon occurred on the same day. Ceiling mounted camera (GoPro Hero4 or Akaso EK7000) recorded the pigeons during these trials. From footage, the duration spent in each of the three zones, as well as the duration spent on perches in each zone, was recorded using BORIS (Friard and Gamba 2016).

### Mass Manipulation

The relationship between body mass and structural size was determined for each of the 2 lofts (see Supplementary Figure 1). The resultant trend line was used to artificially mass load pigeons, by using the residuals from the relationship between size and mass (see Supplementary Fig. 1). Birds who fell under the line of best fit were selected to have artificial mass administered. The resultant negative residual for each pigeon was rounded up to the nearest 5g and that amount was added to each pigeon via the addition of bike balance weights (Qiilu Wheel Weight Balance Self Adhesive, Esonline, Chiswick, London) (Portugal *et al*. 2020), via the back mounted Velcro used to label the pigeons. After the weights had been applied, the same protocol as for the control dominance trials was followed over 10 sessions, in April 2021.

### Plumage Manipulation

Dominance rankings of all the pigeons in each loft were obtained after analysis of the first initial control trials (*n =*10) using David’s score (Gammel *et al*. 2003). The top 20% ranking birds (loft 1 *n =* 4, loft 2 *n =*4) from each loft had all their iridescent neck plumage (which were identifiable due to their colour shifting nature from pink to green and their reflection under UV) covered with black non-toxic acrylic Sharpie markers. These pigeons were habituated to this painting procedure daily, starting three days before the dominance trials were conducted. All pigeons were collected and placed into pigeon carriers. The top ranking pigeons were separated from the carriers and then painted with the black markers until the plumage was a uniform black under natural light. A 345nm UV torch (Lepro, Birmingham, UK) was used to confirm that the plumage reflectance had been obscured (Supplementary Figure 2). Once dry, the painted pigeons were placed in the carriers with the rest of the flock and the dominance trial was started, as previously described. Bird were painted before each dominance trial in the plumage manipulation to ensure consistent colouration in each repeat. This protocol was followed over 10 sessions in June 2021.

### Spectrophotometry

Prior to any plumage manipulations, feather cuttings were taken from the neck of the pigeons, using scissors to cut at the base of the quill to ensure the feature was intact. Only iridescent plumage which appeared green under natural light was sampled, from the top of the neck (McGraw 2004). Gloves were worn whenever handling feathers to prevent contamination (Meadows *et al*. 2011), and feathers were stored in airtight bags out of direct sunlight before spectrophotometry took place.

Feathers were taped to matt black paper overlapping to replicate how they may be arranged *in situ*. Feathers reflectance was analysed using Ocean Optics spectrophotometer (Ocean Optics USB2000, Oxford, UK) measuring between 329nm – 1000nm. Prior to taking measurements the device was calibrated using a WS-1 Diffuse reflectance standard both with and without a light source. The reflectance probe and light source were held at a set distance in a reflectance probe holder at a 90-degree angle from the sample (McGraw 2004). Three repeats per feather sample were taken by placing the reflector holder on the sample at random spots on the tip of the iridescent plumage.

### Respirometry

Resting metabolic rate (RMR) measurements were taken once after the initial dominance and personality control experiments had concluded, and these measurements were taken over a 2-week period in April 2021. Food was removed from the pigeons at 16:00 the day before respirometry trials were performed, to ensure pigeons were post absorptive (Mathot & Dingemanse 2015). Starting at 09:00 the morning after food removal, pigeons were taken in groups of 3-4 from the lofts and transported to the laboratory. Per day, 5-7 pigeons had their RMR measured between approximately 09:00 and 17:00. Each pigeon was individually placed in a respirometry chamber (31 × 23 × 23 cm, volume 12 L) in a dark room and allowed to settle before measurements were taken. Oxygen (O_2_) concentration was calibrated to 20.95% to match ambient air (Lighton, 2008). Initial measurements of ambient air lasting approximately 20 minutes were taken before any pigeon entered the chamber. Ambient air was pulled through the chamber at a flow rate of approximately 1700 mL min^-1^, (SS-4 Sub-Sampler Pump, Sable Systems, Las Vegas, USA) making the flush-out rate of the chamber approximately eight minutes. After recordings for each pigeon (40 minutes) had taken place, another 10 minutes of ambient baseline air was taken. During recordings pigeons were kept within their thermoneutral zone (mean 20.45 ± 3.7 °C (s.e.m.), minimum = 18.2 °C, maximum = 23.5 °C across all traces) (Webster and King 1987).

Air leaving the system first passed through a humidity sensor (RH-300, Sable Systems), where water vapour was removed (anhydrous indicating Drierite®, W. A. Hammond Drierite Co. Ltd, Ohio, USA) before entering the CO_2_ analyser (CA-10a, Sable Systems). CO_2_ was then removed (soda lime, Sigma Aldrich, Merck KGaA, Darmstadt, Germany) and any excess water vapour removed also, before the air then passed through an O_2_ and temperature analyser (FC-10a, Sable Systems). Analysers were connected to a laptop via a UI-2 interface and data were logged in Expedata software (Sable Systems, USA).

### Statistical analysis

Statical analysis was completed using R, v. 3.6.1 (R Software, Vienna, Austria, http://www.R-project.org) (2013). David’s score and hierarchy steepness for each set of trials for each loft was calculated using the package ‘steepness’. Steepness of the dominance hierarchies was calculated as described in de Vries *et al*. (1995) using the R package ‘steepness’ (Leiva and de Vries 2011, R Core Team 2016, de Vries, Stevens and Vervaecke 2006).

All relationships between morphological traits were assessed using linear regression. Correlations between David’s score across trial types were calculated using linear regression and Spearman’s Rank, respectively. Repeatability of David’s score across trials was also assessed using ‘rptR’ package (Schielzeth, Stoffel and Nakagawa 2017) to calculate intraclass correlation coefficient, which describes the amount of potential variation caused by a random effect (in this case, individual variation). The significance was ascertained using a likelihood ratio test with the 95% of repeatability was estimated using 10,000 parametric bootstraps. Repeatability of exploration duration per Zone was also calculated in the same manner. Correlation matrices between measured traits were assessed using the package ‘corrr’. If any two traits exhibited a correlation equal to or above 0.75, one trait was removed prior to further analysis. The relationship between non correlated variables was assessed using a generalised linear model. Trait correlation was also examined for the zone duration aspects of the personality trials. The zone correlation assessments resulted in zones two and three being removed from subsequent analyses. Differences in physiological traits (body mass, tarsus length, maximum % reflectance) and David’s score between sexes was assessed using two sample *t*-tests. The number of interactions per repeat for each trial type was collated and compared using nonparametric Kruskal-Wallace tests; pairwise comparisons were utilised to identify where any significance lied.

## RESULTS

### Phenotypes

Body mass (g) and tarsus length (mm) were significantly correlated in both pigeon flocks combined (LM, R^2^=0.34 F1,29 = 14.95, *P<*0.001; Table 1), with males being significantly larger than females (Two sample *t*-test *t=*-2.39, df=29, *P=*0.02). Resting metabolic rate (RMR) was seen to increase with body mass (LM, R^2^=0.23 F1,29 = 8.53, *P<*0.001), but there was no significant difference in RMR between the two sexes (Two sample *t*-test *t=*-1.41, df=29, *P=*0.17).

No significant difference in maximum feather % reflectance was apparent between the sexes when both flocks were combined (Two sample t-test, t = 0.18, df = 29, *P=* 0.86, after outlier removal (Cook’s Distance, *D*) t = 1.35, df = 26, P = 0.19). RMR was significantly correlated with maximum % reflectance (R^2^=0.30 F1,29 = 11.77, *P=*0.002, after outlier removal (Cook’s Distance, *D*), R^2^=0.44 F1,26 = 20.36, *P*>0.0001). RMR was not significantly correlated with exploratory behaviour in the personality trials (See Supplementary Table 1). Due to the significant relationship between body mass and structural size, only structural size was then used to investigate relationships between size and personality; no significant relationship between any aspects of exploration behaviour and structural size were recorded (GLM, Zone one *t=*-1.7, *P=*0.1, Zone one perch, *t=*0.45, *P=*0.66, Zone two perch, *t=*-1.29, *P=*0.21, Zone three perch, *t=*1.09, *P=*0.29, carrier, *t=*-1.42, *P=*0.17).

Exploration across zones on a population level was highly repeatable carrier; *R* = 0.61 ± 0.08 (s.e.m.), 95% CI:0.43,0.74 *P* < 0.0001, Zone one *R* =0.18 ± 0.08 (s.e.m.), 95% CI:0.02,0.33 *P* < 0.01, zone one perch *R* =0.69 ± 0.07 (s.e.m.), 95% CI:0.52,0.80 *P* < 0.0001, zone two perch *R* =0.11 ± 0.07 (s.e.m.), 95% CI:0.07, 0.26 *P* = 0.04, zone three perch *R* =0.22 ± 0.08 (s.e.m.), 95% CI:0.08,0.05 *P* < 0.001).

**Table 1.** Pigeon phenotypes for two pigeon flocks, loft 1 (*n =* 17 pigeons) and loft 2 (*n =* 14 pigeons). Columns show; pigeon ID, sex, tarsus length (mm), body mass (g), David’s score and associated ranks for each behavioural dominance trial, Trial number refers to (1^st^) control trial, prior to any manipulations, (2^nd^) individuals were weighted, (3^rd^) control dominance trials between manipulations, and (4^th^) dominance trial in which the top 20% ranked birds from the 1^st^ control from each loft had their iridescent plumage dampened resulting in decreased reflectance, maximum iridescent feather reflectance (%) for each individual. Loft refers to which flock each bird was part, of either 1 or 2.

| Bird ID | Sex | Tarsus (mm) | Mass (g) | 1st Trial David's score | 1st Trial rank | 2nd Trial David's score | 2nd Trial rank | 3rd Trial David's score | 3rd Trial rank | 4th Trial David's score | 4th Trial rank | Reflectance (%) | RMR ml/min | Loft no. |
| --- | --- | --- | --- | --- | --- | --- | --- | --- | --- | --- | --- | --- | --- | --- |
| 1 | M | 36.33 | 460 | -39.78 | 15 | -19.98 | 13 | -8.09 | 10 | 45.56 | 4 | 72.06 | 10.04 | 1 |
| 2 | F | 33.25 | 460 | -64.32 | 17 | -59.48 | 17 | -80.25 | 17 | -51.07 | 16 | 90.37 | 13.66 | 1 |
| 3 | M | 36.13 | 500 | -27.20 | 12 | -15.16 | 10 | -17.59 | 11 | 29.82 | 6 | 68.43 | 16.03 | 1 |
| 4 | F | 32.80 | 400 | -59.04 | 16 | -13.17 | 9 | -21.73 | 13 | -30.02 | 13 | 105.41 | 7.41 | 1 |
| 5 | F | 33.34 | 500 | -13.11 | 10 | 9.66 | 6 | 22.70 | 5 | 22.49 | 7 | 120.27 | 8.45 | 1 |
| 6 | M | 35.85 | 470 | 58.82 | 2 | 59.07 | 2 | 50.71 | 3 | 47.10 | 3 | 79.30 | 14.71 | 1 |
| 7 | F | 34.37 | 430 | 38.14 | 4 | -21.25 | 15 | -19.28 | 12 | -29.93 | 12 | 108.06 | 8.29 | 1 |
| 8 | M | 37.43 | 520 | 51.97 | 3 | 62.33 | 1 | 69.10 | 1 | 49.18 | 2 | 57.40 | 15.77 | 1 |
| 9 | M | 35.74 | 440 | 73.16 | 1 | 38.79 | 3 | 50.98 | 2 | 35.90 | 5 | 75.85 | 15.63 | 1 |
| 10 | F | 34.64 | 420 | -29.44 | 13 | -38.27 | 16 | -61.48 | 16 | -53.66 | 17 | 99.26 | 7.86 | 1 |
| 11 | F | 34.53 | 460 | -15.63 | 11 | -15.67 | 11 | 2.26 | 8 | -43.43 | 15 | 77.41 | 8.99 | 1 |
| 12 | M | 36.26 | 480 | 22.11 | 6 | 35.11 | 4 | 16.77 | 6 | 52.81 | 1 | 83.30 | 10.74 | 1 |
| 13 | M | 34.15 | 450 | -9.47 | 9 | 9.32 | 7 | -26.87 | 14 | -17.93 | 11 | 78.37 | 8.52 | 1 |
| 14 | M | 36.78 | 450 | 35.27 | 5 | -8.20 | 8 | -31.16 | 15 | -10.76 | 9 | 47.29 | 15.12 | 1 |
| 15 | F | 32.32 | 410 | -35.46 | 14 | -21.17 | 14 | 9.77 | 7 | -32.48 | 14 | 143.75 | 9.23 | 1 |
| 16 | F | 33.64 | 480 | 20.50 | 7 | 14.63 | 5 | 47.25 | 4 | -14.56 | 10 | 84.08 | 11.65 | 1 |
| 17 | M | 36.91 | 500 | -6.50 | 8 | -16.57 | 12 | -3.10 | 9 | 0.98 | 8 | 80.36 | 8.40 | 1 |
| A | M | 36.02 | 510 | 10.86 | 8 | 10.35 | 5 | 11.03 | 5 | 23.40 | 4 | 130.71 | 8.58 | 2 |
| B | F | 34.25 | 420 | -48.35 | 13 | -15.59 | 12 | -15.48 | 11 | 13.22 | 7 | 78.91 | 12.30 | 2 |
| C | F | 36.32 | 450 | -47.72 | 12 | -13.92 | 11 | -16.89 | 12 | -21.42 | 11 | 20.36 | 10.16 | 2 |
| D | M | 35.44 | 490 | 4.46 | 9 | 29.22 | 2 | 36.96 | 2 | 24.77 | 3 | 99.74 | 10.80 | 2 |
| E | F | 34.70 | 530 | 28.18 | 4 | 21.85 | 3 | 25.48 | 3 | 20.23 | 5 | 73.07 | 16.32 | 2 |
| F | M | 34.80 | 470 | 29.38 | 3 | 19.22 | 4 | 23.01 | 4 | 35.29 | 1 | 80.31 | 13.82 | 2 |
| G | F | 33.21 | 410 | 23.48 | 5 | 3.15 | 6 | 3.99 | 6 | 2.38 | 8 | 83.69 | 12.41 | 2 |
| H | F | 31.86 | 440 | -62.02 | 14 | -42.28 | 14 | -53.65 | 14 | -42.23 | 14 | 65.03 | 8.91 | 2 |
| I | M | 36.70 | 560 | 48.56 | 1 | 49.25 | 1 | 52.42 | 1 | 30.47 | 2 | 52.39 | 17.74 | 2 |
| J | M | 35.12 | 500 | 36.46 | 2 | -0.86 | 7 | -0.38 | 8 | 14.41 | 6 | 105.70 | 9.19 | 2 |
| K | F | 33.34 | 460 | -21.26 | 10 | -6.68 | 9 | -11.19 | 9 | -12.43 | 10 | 103.07 | 10.78 | 2 |
| L | F | 35.74 | 440 | 17.69 | 7 | -11.67 | 10 | -15.01 | 10 | -36.00 | 12 | 45.32 | 15.44 | 2 |
| M | M | 33.17 | 380 | 21.73 | 6 | -3.61 | 8 | 1.60 | 7 | -11.51 | 9 | 146.05 | 7.01 | 2 |
| N | F | 32.50 | 420 | -41.47 | 11 | -38.44 | 13 | -41.89 | 13 | -40.58 | 13 | 71.61 | 6.43 | 2 |

### Dominance

Generalised linear models for each separate flock showed no significant relationships between David’s score and any covariates (tarsus length, maximum % reflectance, resting metabolic rate, duration in zone one, zone one perch, zone two perch, zone three perch and carrier). When data for both flocks were combined, David’s score was best predicted by tarsus length (GLM, *t=* 2.17, *P=*0.04), maximum % reflectance (GLM, *t=* 2.41, *P=*0.02), and resting metabolic rate (GLM, *t=*2.51, *P=* 0.02 (Table 2). The relationship between RMR and David’s score remained significant after accounting for body mass (GLM, *t=*2.53, *P=* 0.02). On average, males occupied higher ranking positions than females (Two sample *t*-test t =-3.3 df = 29, p-value = 0.0002). The steepness of the dominance hierarchies remained similar throughout the dominance trials (see Supplementary Table 2).

**Figure 1.**
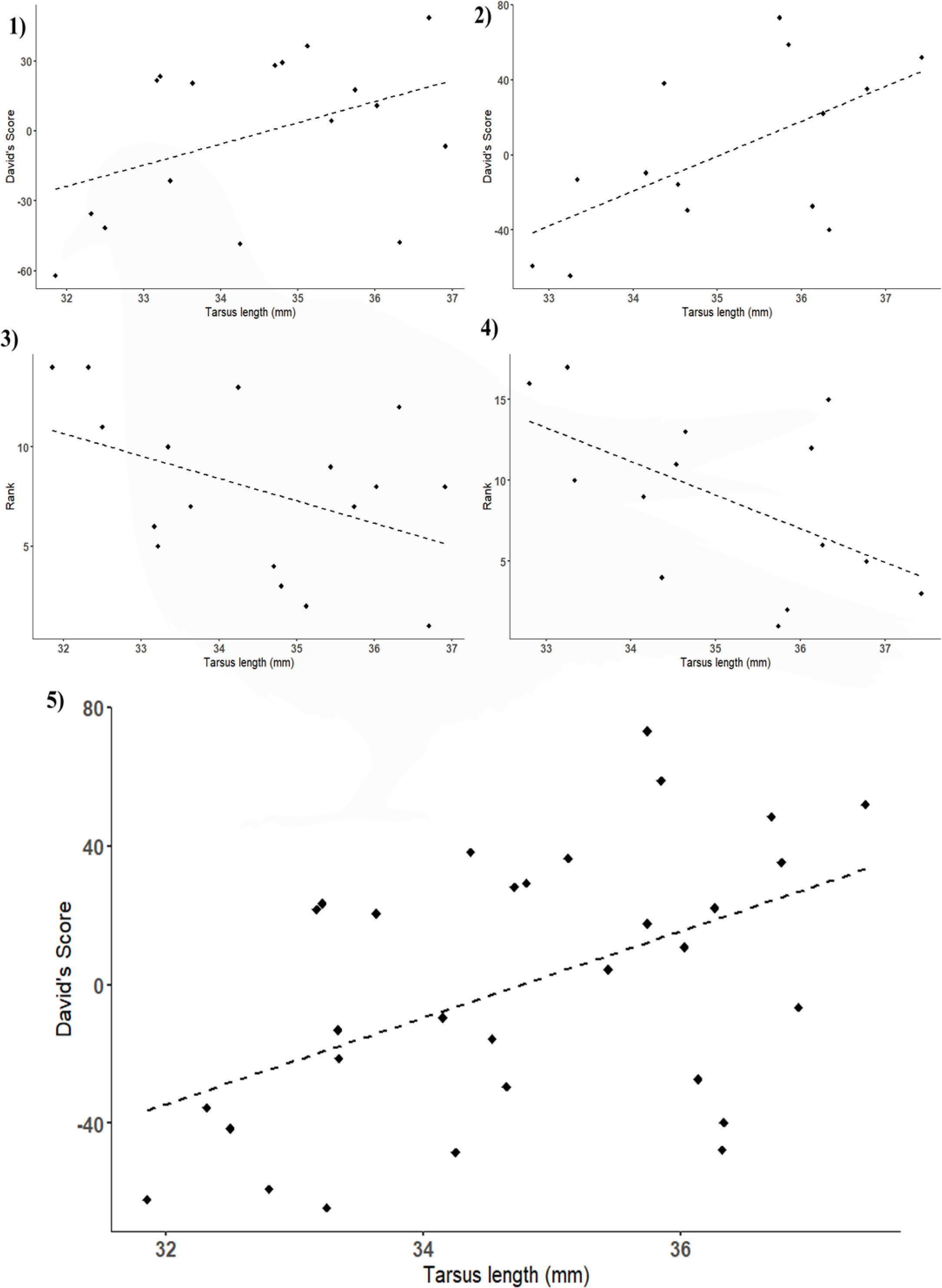
1-2) Relationship between David’s score and Tarsus length (F1,15 = 5.91, *P=*0.03 R^2^=0.28), and rank and tarsus length (F1,15 = 6.19, *P=*0.03, R^2^=0.29) respectively for loft 1 (*n=* 17 pigeons). 3-4) Relationship between David’s score and tarsus length (F1,12 = 3.11, *P=*0.1, R^2^=0.21) and rank and tarsus length (F1,12 = 2.39, *P=*0.15, R^2^=0.17), respectively, for loft 2 (*n=* 14 pigeons). 5) presents data for both lofts combined for David’s score (F1,29 = 7.41, *P=*0.01, R^2^=0.2) All *P*-values are at a 95% confidence level.

**Table 2.** Generalised linear model results for both lofts combined (31 pigeons) between initial David’s score (dominance) as a response variable and structural size (tarsus length), maximum reflectance% = maximum reflectance % of iridescent feathers, resting metabolic rate (RMR), and exploratory behaviour (zone one, zone one perch, zone two perch, zone three perch and carrier) (see *Methods*) as explanatory variables. All *P*-values are at a 95% confidence level. Bold denotes significant.

|  | Estimate | Std.<br>Error | *t*-value | p |
| --- | --- | --- | --- | --- |
| Intercept | -514.10 | 185.60 | -2.77 | 0.01 |
| Tarsus length (mm) | 11.13 | 5.14 | 2.17 | **0.04** |
| Maximum %<br>reflectance | 0.680 | 0.281 | 2.41 | **0.02** |
| Zone one | 0.001 | 0.17 | 0.007 | 0.99 |
| Zone one perch | 0.02 | 0.02 | 0.96 | 0.35 |
| Zone two perch | -0.01 | 0.02 | -0.75 | 0.46 |
| Zone three perch | 0.002 | 0.004 | 0.38 | 0.71 |
| Carrier | 0.004 | 0.02 | 0.26 | 0.80 |
| RMR | 6.00 | 2.39 | 2.51 | **0.02** |

### Rank stability

Dominance rank and David’s score remained stable across the two control trials and over the course of the two manipulation dominance trials (See Supplementary Fig. 2), with David’s score and rank being highly correlated between trials in both flocks, throughout (see Table 5, Table 6, for full Spearman’s rho and *P* values). In addition, David’s score across all trials for both flocks were highly repeatable (loft 2, *R* =0.79 ± 0.09 (s.e.m.), 95% CI:0.56,0.90 *P* < 0.0001; loft 2, *R* =0.71± 0.10 (s.e.m.), 95% CI:0.45,0.84, *P* < 0.0001).

Aggression levels between trial types varied for both flocks (loft 1 Kruskal-Wallis X^2^ = 10.99, df = 3, p-value = 0.012; loft 2 Kruskal-Wallis X^2^= 10.38, df = 3, *P*-value = 0.016) (Table 5). Pairwise comparison indicated this significance was between the 1^st^ control and mass manipulation trial (*P=*0.03), and the mass manipulation and feather manipulation trials (*P=*0.006) in loft 2, and between the 1^st^ control and mass manipulation trial (*P=*0.04) in loft 1. Social network diagrams for each trial type (Fig. 3) show similar interactions between individuals in each trial type. Though interactions in the 2^nd^ trial (mass manipulation) for both lofts show a shift in dynamic, with a large proportion of interactions during these trials taking place between the same 5-6 birds.

**Table 3.** Spearman’s rank (rho) (italicized and shaded) comparisons of ranks for both lofts between control dominance trials and trials with manipulated mass or plumage. *P* values (non-italicized) between trials <0.01 suggest correlation between ranks between trials. Trial number refers to (1^st^) control trial, prior to any manipulations, (2^nd^) specific individuals were weighted, (3^rd^) control dominance trials between manipulations, and (4^th^) dominance trial in which the top 20% ranked birds from the 1^st^ control from each loft had their iridescent plumage dampened resulting in decreased reflectance.

| Loft 1 | 1<sup>st</sup><br>Trial | 2<sup>nd</sup><br>Trial | 3<sup>rd</sup><br>Trial | 4<sup>th</sup><br>Trial | Loft 2 | 1<sup>st</sup><br>Trial | 2<sup>nd</sup><br>Trial | 3<sup>rd</sup> Trial | 4<sup>th</sup> Trial |
| --- | --- | --- | --- | --- | --- | --- | --- | --- | --- |
| 1<sup>st</sup> Trial | **X** | >0.001 | 0.01 | 0.02 | 1<sup>st</sup> Trial | **X** | 0.002 | 0.003 | 0.01 |
| 2<sup>nd</sup> Trial | 0.70 | **X** | >0.001 | >0.001 | 2<sup>nd</sup> Trial | 0.77 | **X** | >0.001 | >0.001 |
| 3<sup>rd</sup> Trial | 0.59 | 0.75 | **X** | 0.01 | 3<sup>rd</sup> Trial | 0.75 | 0.99 | **X** | >0.001 |
| 4<sup>th</sup> Trial | 0.58 | 0.74 | 0.65 | **X** | 4<sup>th</sup> Trial | 0.65 | 0.89 | 0.89 | **X** |

**Table 4.** Linear regressions comparing David’s score between control dominance trials and manipulated mass or plumage trials for both lofts. Trial number refers to (1^st^) control trial, prior to any manipulations, (2^nd^) specific individuals were weighted, (3^rd^) control dominance trials between manipulations, and (4^th^) dominance trial in which the top 20% ranked birds from the 1^st^ control from each loft had their iridescent plumage dampened resulting in decreased reflectance. All trials were significantly correlated with each other trial, suggesting rank stability.

| David's<br>score<br>comparison<br>Loft 1 | F Degrees of<br>freedom | R<sup>2</sup> | P | David's<br>score<br>comparison<br>Loft 2 | F Degrees of<br>freedom | R<sup>2</sup> | P |
| --- | --- | --- | --- | --- | --- | --- | --- |
| 1<sup>st</sup> trial vs<br>2<sup>nd</sup> trial | F<sub>1,15</sub>=23.5 | 0.61 | *P*>0.001 | 1<sup>st</sup> trial vs<br>2<sup>nd</sup> trial | F<sub>1,12</sub>=19.19 | 0.62 | *P*>0.001 |
| 1<sup>st</sup> trial vs<br>3<sup>rd</sup> trial | F<sub>1,15</sub>=13.2 | 0.47 | *P*=0.002 | 1<sup>st</sup> trial vs<br>3<sup>rd</sup> trial | F<sub>1,12</sub>=20.06 | 0.63 | *P*>0.001 |
| 1<sup>st</sup> trial vs<br>4<sup>th</sup> trial | F<sub>1,15</sub>=5.4 | 0.26 | *P*=0.035 | 1<sup>st</sup> trial vs<br>4<sup>th</sup> trial | F<sub>1,12</sub>=7.702 | 0.40 | *P*=0.017 |
| 2<sup>nd</sup> trial vs<br>3<sup>rd</sup> trial | F<sub>1,15</sub>=47.8 | 0.76 | *P*>0.0001 | 2<sup>nd</sup> trial vs<br>3<sup>rd</sup> trial | F<sub>1,12</sub>=1116.0 | 0.99 | *P*>0.0001 |
| 2<sup>nd</sup> trial vs<br>4<sup>th</sup> trial | F<sub>1,15</sub>=18.9 | 0.56 | *P*=0.001 | 2<sup>nd</sup> trial vs<br>4<sup>th</sup> trial | F<sub>1,12</sub>=32.8 | 0.73 | *P*>0.0001 |
| 3<sup>rd</sup> trial vs<br>4<sup>th</sup> trial | F<sub>1,15</sub>=12.2 | 0.45 | *P*=0.003 | 3<sup>rd</sup> trial vs<br>4<sup>th</sup> trial | F<sub>1,12</sub>=36.3 | 0.75 | *P*>0.0001 |

**Figure 2.**
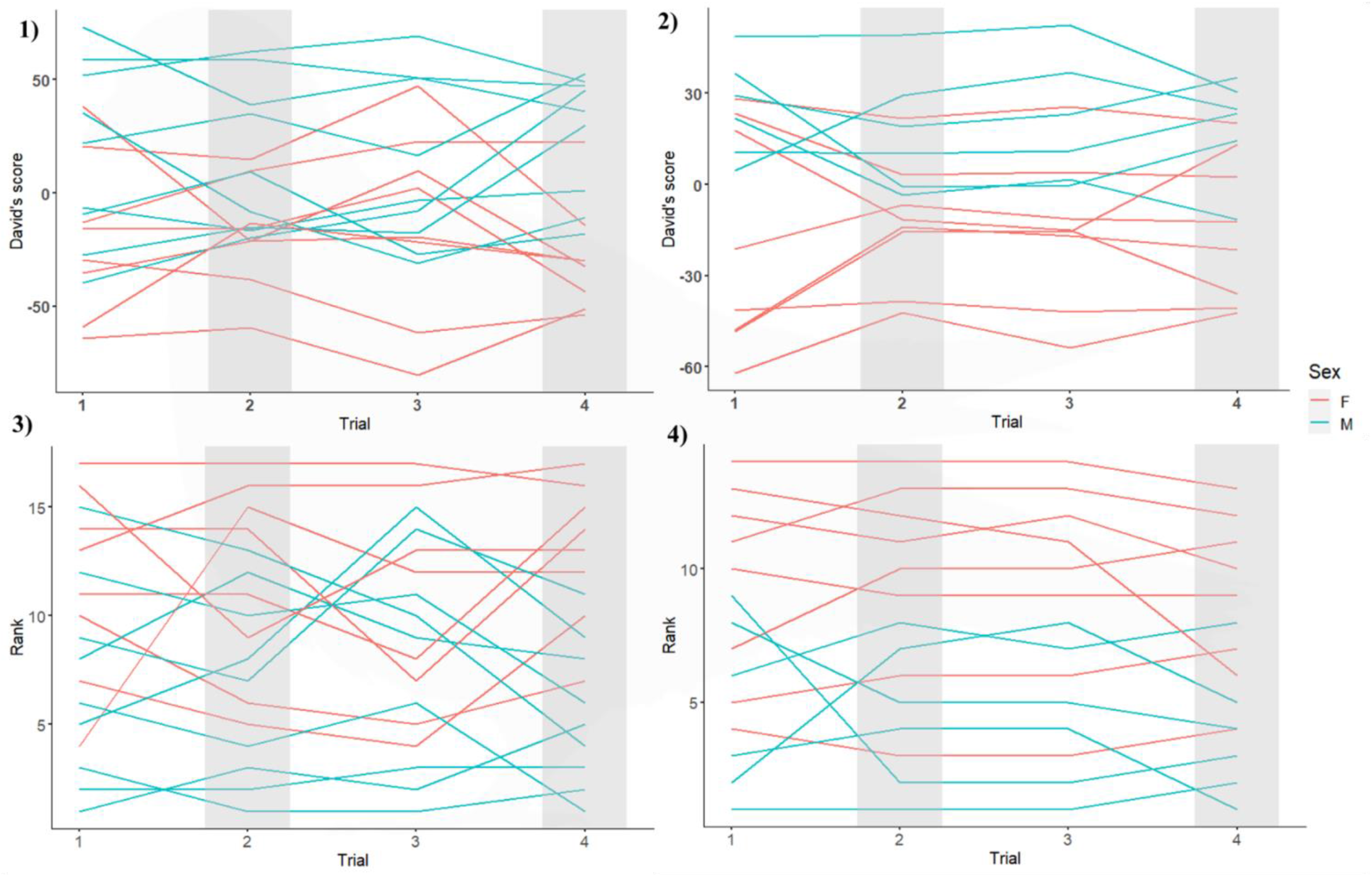
David’s score and rank over the two control trials (1 and 3 denoting the 1^st^ and 2^nd^ control respectively on x-axis) for two flocks of pigeons (loft 1, *n=*17; loft 2, *n =* 14). The mass manipulation (3 on x-axis) and plumage manipulation (4 on x-axis) re denoted by grey shaded area. 1-2 show results for loft 1 and 3-4 show results for loft 2. Colours represent male and female pigeons (see legend).

**Table 5.**
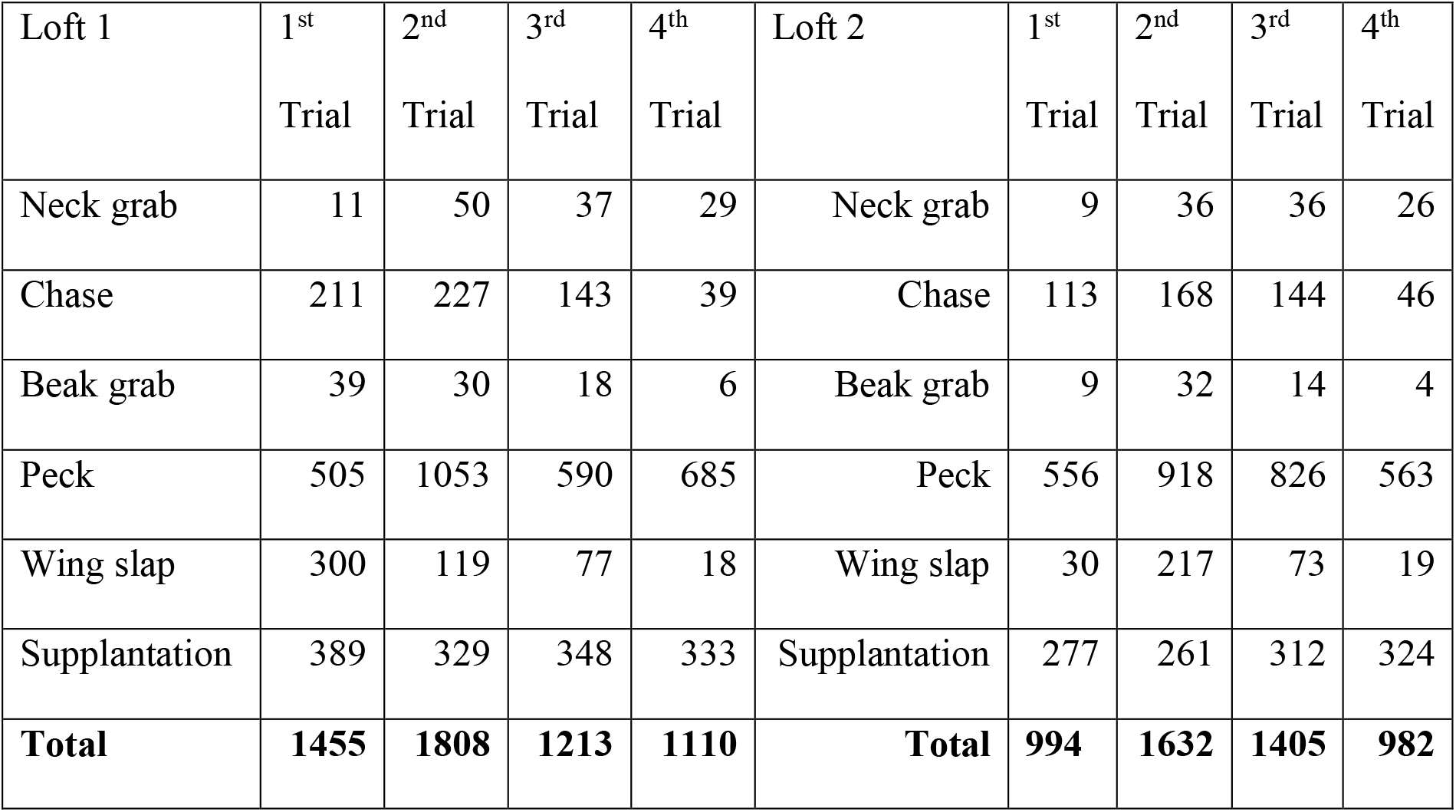
Composition of aggressive behaviours across 4 dominance trials in two flocks of pigeons (see methods for definition of each behaviour type). Trial number refers to (1^st^) control trial, prior to any manipulations, (2^nd^) individuals were weighted, (3^rd^) control dominance trials between manipulations, and (4^th^) dominance trial in which the top 20% ranked birds from the 1^st^ control from each loft had their iridescent plumage dampened resulting in decreased reflectance.

**Figure 3.**
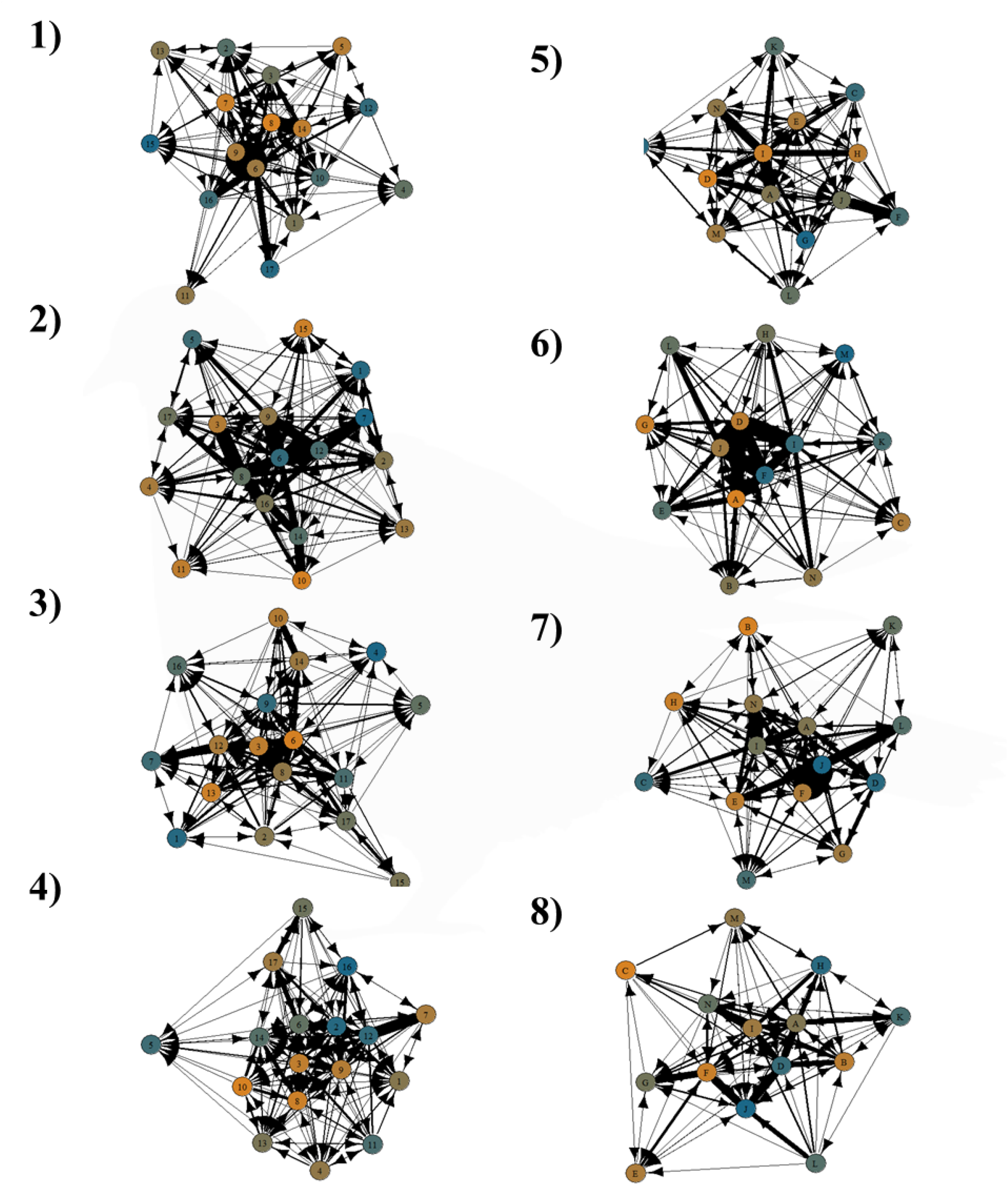
Social network of aggressive interactions for both control (1^st^ control 1 and 5, 2^nd^ control 3 and 7 and manipulated trials (mass manipulation 2 and 6, plumage manipulation 4 and 8) for loft 1 (*n* = 17 pigeons) diagrams 1-4 and for loft 2 (*n* = 14 pigeons) diagrams 5-8. Arrows display aggressive interactions between two individuals, with the direction and thickness indicating the number and recipient of the interaction. Nodes are coloured by social status, from dominant (orange) to subordinate (blue). Letters (loft 2) and numbers (loft 1) refer to a single pigeon; see Table 1 for labels and pigeon IDs.

## DISCUSSION

Over the four dominance trial types conducted, the hierarchies for both pigeon flocks remained stable across all experimental conditions, and where not destabilised by any experimental perturbations. Analysis of morphological traits revealed a clear positive relationship between aggressive behaviour (via David’s score) and body size, body mass and resting metabolic rate (RMR). Additionally, there was a negative relationship between RMR and iridescent plumage % reflectance.

### Intrinsic Traits and David’s score

Previous research on homing pigeons (Portugal *et al*. 2017a; 2017b) reported a strong relationship between body mass and dominance rank. This is prevalent across multiple taxa, potentially indicating that larger body sizes and masses – the two typically being correlated – are important factors for resource holding, likely due to the increased ability to exclude and intimidate smaller individuals if you are bigger and heavier (Brown and Maurer 1986; Márquez-Luna *et al*. 2019). Such a scenario likely contributes to a positive feedback loop, as the already heavier, more dominant, birds obtain a larger quantity of resources, which in turn assists in body mass maintenance and mass gains.

Resting metabolic rate predicted aggressive behaviour (David’s score), even when body mass was accounted for which is in accordance with the *performance model* (Mathot, Dingemanse and Nakagawa 2019). As previously described, the *performance model* suggests individuals with higher RMRs perform a higher amount of energetically costly behaviours, in addition to behaviours which result in net energy gain (Mathot, Dingemanse and Nakagawa 2019). In contrast to the relationship with David’s score, RMR was not predictive of an explorative personality, which directly contradicts the aforementioned *performance model*. In this case, the results suggests that investment in aggression (leading to higher David’s score) contributed to a net gain of energy due to better access to food resources (Careau *et al*. 2008).

No relationship between any of the recorded personality traits and David’s score were observed. Similar studies have reported a variety of contrasting and corroborating results. Some studies indicate a relationship between heightened boldness and social dominance rank, such as negative relationships in captive black cap chickadees *Poecile atricapillus* with boldness being a trait more prevalent in subordinate birds (MacDougall-Shackleton *et al*. 2011). In contrast, other species and populations exhibit no relationship between rank and personality, such as that seen in wild black cap chickadees (Devost *et al*. 2016) and common waxbills *Estrilda astrild* (Funghi *et al*. 2014).

### Personality and RMR

In addition to the lack of relationship between personality and David’s score, a similar lack of relationship was observed between resting metabolic rate and personality traits. Careau and Garland (2012), in addition to Mathot and Dingemanse (2015), reviewed empirical research which found relatively few correlations of boldness or exploration with metabolic rate in avian species. Two studies in great tits have indicated negative correlations between risk taking behaviour and basal metabolic rate (BMR) (Mathot *et al*. 2014), in addition to exploration and BMR specifically in females (Bouwhuis *et al*. 2014). Both of these studies were conducted with wild populations where individuals are subject to different selection and resource pressures. In contrast, the homing pigeons in the present study are domesticated and kept in captivity, where access to food and water were unlimited, and the birds were safe from being predated upon when in their home loft. This difference in overall selection pressures could have impacted general motivation and willingness to explore during trials (Montiglio *et al*. 2018, Poranen, and Ruuskanen 2021). An interesting further avenue of study would be to repeat the experiments with wild feral pigeons, to determine if they match the behaviours of the captive pigeons. In the present study, pigeons explored an unfamiliar loft alone. It is possible that in the case of homing pigeons, navigation and flight exploration could be a more reliable indicator of boldness (Portugal *et al*. 2017a), as opposed to ground-based experiments. Group flight is an energetically costly activity (Taylor *et al*. 2019; Sankey *et al*. 2019) and captive homing pigeons will experience similar predation threats during homing flights that wild populations encounter. Under these circumstances, exploratory flight and RMR may be linked in homing pigeons to a greater extent than loft based explorations.

Previous work establishing the links between metabolic rate, dominance and individual personality traits have not been ubiquitous. In the present study, it is possible that time of day was potentially impacting personality. Personality trials took place throughout the day between 09:00 and 17:00, and the time of day at which individuals had their personalities assessed could have been influenced by their natural behavioural circadian rhythms. Typically, pigeons are less active in the afternoons (Ricketts *et al*. 2021). Thus, birds who had repeat trials in the afternoons may exhibited reduced activity levels and a lowered willingness to explore. Individuals may have also been influenced by the external temperatures; personality assays were undertaken in February and March 2021, where temperatures averaged 8◦C and ranged from 16◦C to -2 ◦C. These low temperatures may have affected motivation for exploration leading to variation in activity (see Paladino 1995).

A consistent issue raised by prior studies is that of measuring personality in a social species, with isolated individuals (Funghi *et al*. 2014). Several studies on zebra finches indicate that individuals exhibit decreased neophobia when solitary, compared to social contexts which can appear counter intuitive (Schuett and Dall, 2009, Mainwaring, Beal and Hartley 2011, Kerman, Miller and Sewall 2018). Zebra finches with higher basal metabolic rates were also more likely to adopt the scrounger role as a foraging strategy in groups (Mathot *et al*. 2009). As personalities of individuals of social species can be so different when isolated in contrast to within a group, a more accurate measure of personality may be to assess neophobic tendencies of individuals during both solitary and group settings. This method may reveal more about individual behavioural syndrome and social standing (Funghi *et al*. 2014).

### Iridescent feather % reflectance and resting metabolic rate

Maximum % reflectance of iridescent neck feathers was seen to decrease with increasing RMR. This suggests a potential energetic cost of either producing or maintaining the brightness of iridescent plumage. On a genetic level, sequencing of both iridescent and non-iridescent plumage related genes of superb starlings (*Lamprotornis superbus*) indicated non-iridescent feathers are associated with higher metabolic gene expression compared to iridescent plumage (Rubenstien *et al*. 2021). These different plumage types not only differ in production but also greatly differ in maintenance. A study on the iridescent plumage of mallard ducks *Anas platyrhynchos* indicated iridescent feathers were less hydrophobic and less efficient at self-cleaning, due to the difference in barbule structure (Eliason and Shawkey 2011). This functional difference could suggest that individuals have to invest a greater amount of time preening iridescent feathers to maintain structurally coloured plumage, which is likely to come at an energetic cost (Viblanc *et al*. 2011).

The negative relationship between (assumed) feather quality and David’s score has been reported in other species. Dominant coal tits *Periparus ater* produced lower quality feathers that grew at a slower rate than subordinates, thought to be a trade off with the costs of increased territory size, resources and mate choice (Hay *et al*. 2004).

### Iridescent feather %reflectance as a dominance signal

Though GLM results indicated a significant relationship between maximum % reflectance of plumage and David’s score, this does not necessarily suggest that feather reflectance is a dominance signal in homing pigeons, as no significant change in dominance rank occurred when feathers and associated plumage were altered. The lack of significant overall aggressive interactions towards plumage altered birds suggests birds were still recognising each other despite the painting of certain feathers. This is corroborated by studies which have demonstrated that individual recognition in pigeons is more focused on their ‘face’ than their body (Shimizu 1998). Furthermore, the observed relationship between dominance position and reflectance was negative, which would be unusual for a dominance signal; dominance signals are commonly highly investing ornamental traits.

While in the present study altering plumage signals had no significant effect on positioning within the hierarchy, dampening experiments on blue tits did show that reflectance was used to gauge attackers rank, but only during initial encounters. Once the birds were familiar with each other, subsequent interactions ended the same as the first confrontation, regardless of any reversal in UV plumage alterations (Vedder *et al*. 2010). This reliance on first interactions for determining dominance could suggest that because the pigeons in the present study had been living together for many years, the alteration of this signal were ineffective in disrupting hierarchies. Plumage brightness in house finches *Carpodacus mexicanus* was negatively associated with dominance, with drab males being more aggressive and having poorer access to food. McGraw and Hill (2000) suggested this could be due to birds having to attack more to make up for their poor plumage and diet to obtain resources. Dominant individuals usually initiate more aggressive interactions and retreat less than subordinates, spending a proportionally larger time in combat (Hay *et al*. 2004). These birds also predominately have higher activity levels (Ricketts *et al*. 2021) and this could be attributed to their higher metabolic costs.

### Hierarchy stability

Between all dominance hierarchy trials conducted there was no significant change in rank, with neither attempts to perturb hierarchy organisation being successful. This suggests that pigeon dominance hierarchies are highly stable over time and resistance to major disruption. This may particularly be the case in larger groups of more than 10 individuals, as previously observed in carrion crows (Chiarati *et al*. 2010) and Lake Tanganyika cichlids *Neolamprologus pulcher* (Taborsky *et al*. 2005)

Artificially increasing mass of lighter individuals did not significantly affect hierarchy structure, which could indicate perhaps that body size is a more important trait for asserting dominance than mass (Márquez-Luna *et al*. 2019, Brown and Maurer, 1986). Given that homing pigeons have context dependant hierarchies (Nagy *et al*. 2013), this relationship between size, RMR and rank and the associated rank stability is likely different when navigating during flight. Watts *et al*. (2016) indicated that leaders during flight lose influence over flock mates when they have inaccurate information. In those scenarios the consequence for spreading misinformation is much higher, compared to on the ground. The fact that the ground-based hierarchy was able to remain stable despite perturbations suggests none of the altered traits were manipulated sufficiently, or the traits that were focused upon don’t contribute to hierarchy stability. Given that hierarchy formation reduces group aggression, it is understandable why hierarchies remain stable.

Most empirical evidence suggests dominance hierarchies are usually highly stable, with reported instability only occurring when there is a change in group composition through emigration or immigration (Grossel *et al*. 2021). In the present study, the two populations remained stable except for loft 2 where, due to an unrelated incident, two pigeons were removed. While the overall flock hierarchy structure did not significantly change as a result of this, there is some evidence to suggest there was some instability caused by this change with one or two members changing more than 2 ranking positions. Whether this was caused by the flock changes or just seasonal variation is unknown, as similar occurrences were visible in loft 1, which in contrast had a stable undisturbed population over all 40 dominance trials.

## Conclusion

Overall, our results indicate dominance rank in homing pigeons is positively correlated with tarsus length and resting metabolic rate. In addition, a negative relationship was observed between iridescent plumage reflectance and David’s score, suggesting a link to the further cost of dominance traded off against the high investment costs of iridescence. The dominance hierarchies of the homing pigeons were highly resistance to experimental perturbations, indicating how their formation in social species is highly beneficial through decreasing aggression levels and maintaining stable group dynamics.

We thank the following people for useful discussions: Marie Attard, Elli Leadbeater, David Morritt. We are grateful to Stephanie McClelland and Cecylia Watrobska for help with looking after the birds. Funding was provided by a Royal Society Research Grant (R10952) to S.J.P. All experiment protocols were approved by the RHUL local Ethics and Welfare Committee

## AUTHOR CONTRIBUTIONS

Amaia A. Urquia-Samele: Conceptualization (equal); Data curation (lead); Formal analysis (lead); Investigation (lead); Methodology (equal); Writing-original draft (lead); Writing-review & editing (equal).

Steven J. Portugal: Conceptualization (equal); Funding acquisition (lead); Investigation (supporting); Methodology (equal); Project administration (lead); Supervision (lead); Writing original draft (supporting); Writing-review & editing (equal).

## COMPETING INTERESTS

We declare we have no competing interests.

### Data availability statement

All data are available within the tables provided in the main text and supplementary information.

