## Supplementary Material for "Homing pigeon (*Columba livia*) dominance hierarchies are stable over time and resistant to perturbation"

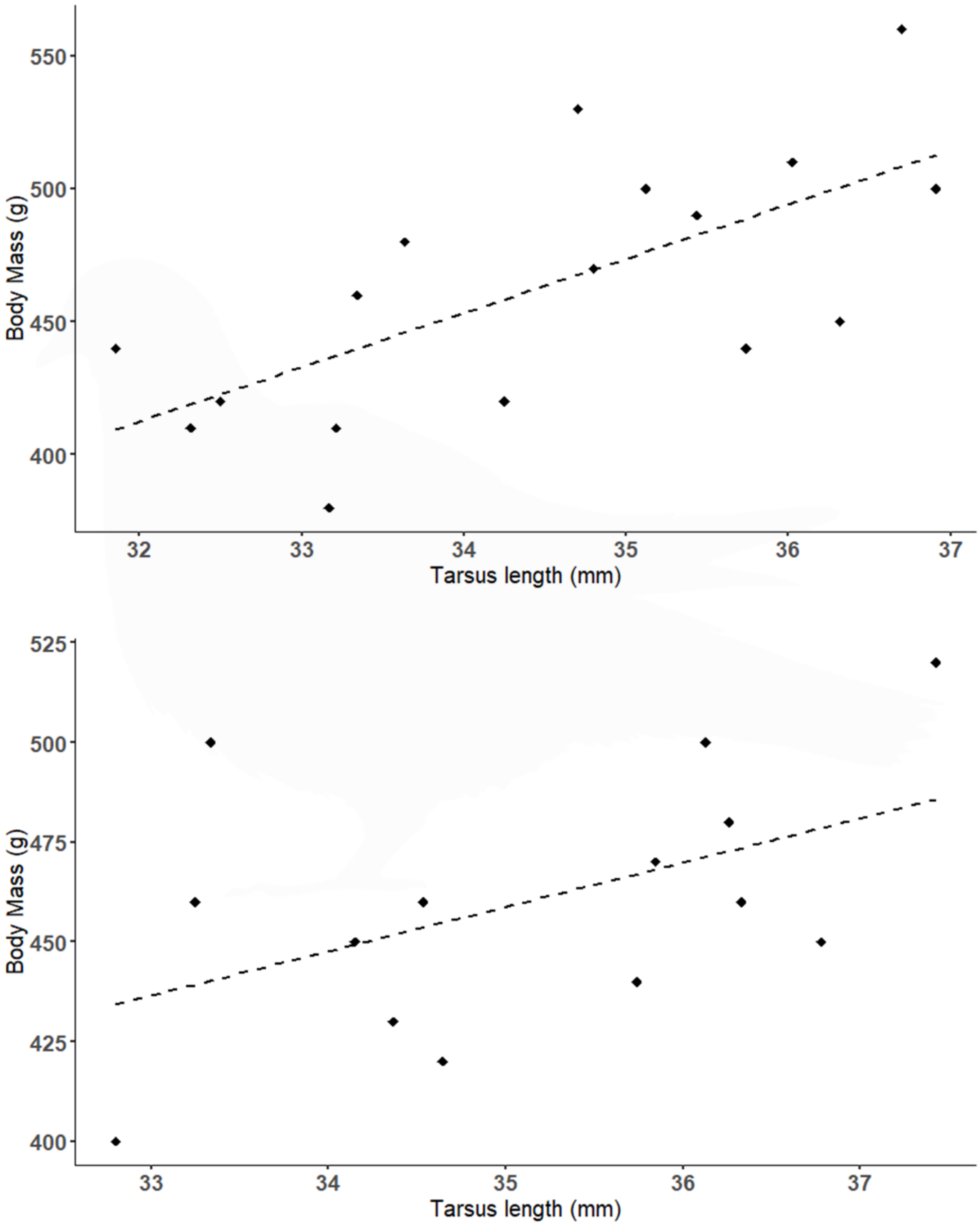


**Supplementary Figure 1.** Body mass and tarsus length for both loft 1 (1) and loft 2 (2), all individuals that fell below the line of best fit had their body mass artificially increased using the equation of the line, (loft 1, y=23.01x-333.33, $R^2$= 0.32, *P*=0.2, loft 2, y=17.58x-162.22, $R^2$= 0.42, *P*=0.1).

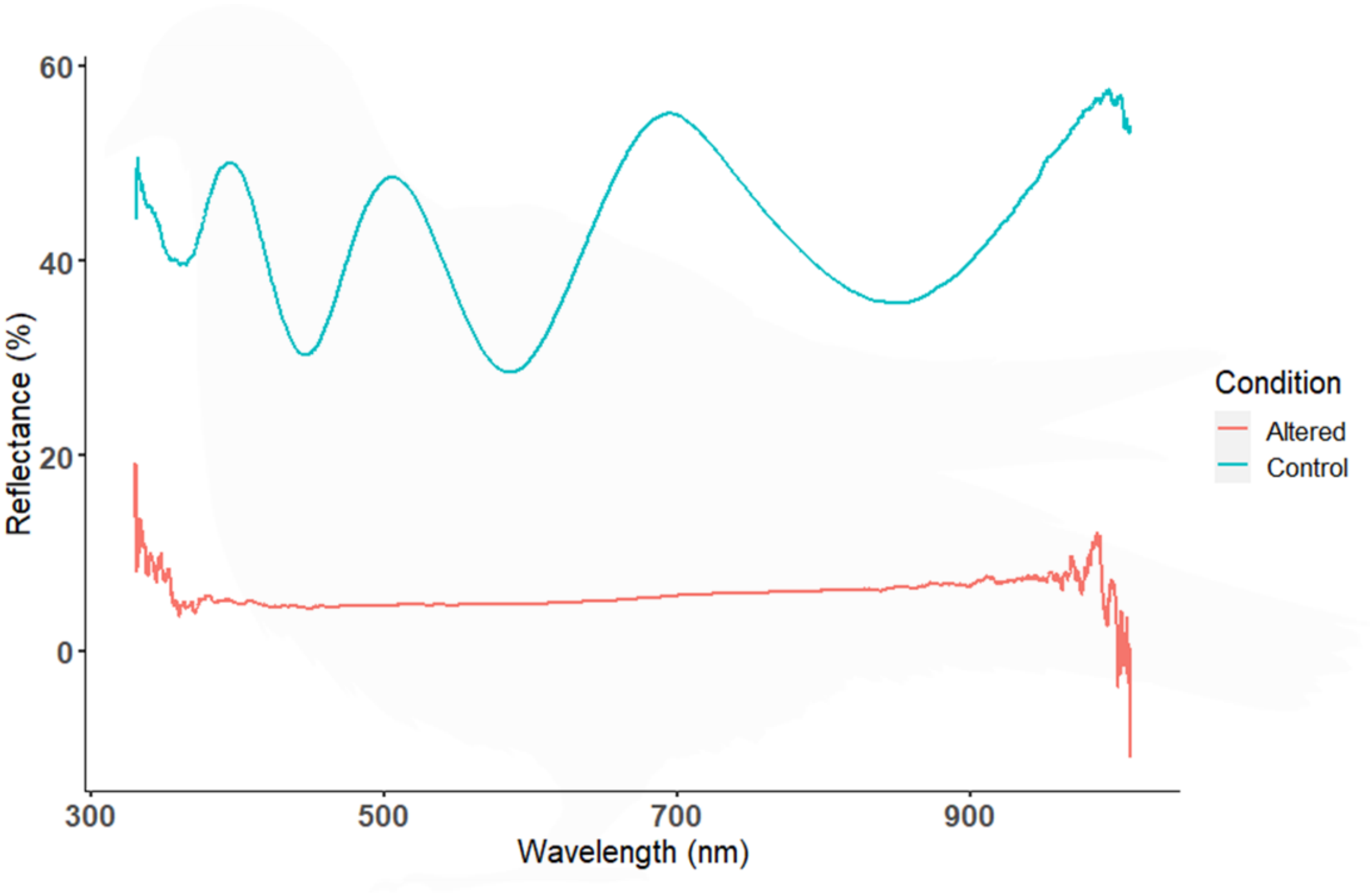


**Supplementary Figure 2.** Mean reflectance spectra of top 20% of ranking birds for loft 1(*N* =4) and loft 2 (*N* =3) with no manipulation (control, blue line). Overall manipulated birds had their iridescent plumage reflectance reduced by 71% on average.

**Supplementary Table 1.** Generalised linear model results between resting metabolic rate (RMR) as response variable and exploratory behaviour described by the duration spent in each zone (zone one, zone one perch, zone two perch, zone three perch, carrier; see *Methods*) as explanatory variables. All *P*-values are at a 95% confidence level.

| | Estimate | Std. Error | *t*-value | P |
|---|---|---|---|---|
| Intercept | 11.98 | 0.92 | 12.96 | >0.0001 |
| Zone one | -0.0085 | 0.02 | -0.52 | 0.61 |
| Zone one perch | <0.001 | 0.001 | 0.32 | 0.75 |
| Zone two perch | 0.001 | 0.002 | -0.92 | 0.37 |
| Zone three perch | <0.001 | <0.001 | 0.68 | 0.5 |
| Carrier | -0.002 | 0.001 | -1.36 | 0.19 |

**Supplementary Table 2.** Steepness hierarchy parameters for loft 2 (*n* = 17 pigeons) and loft 3 (*n* = 14 pigeons). Trial number refers to ($1^{st}$) control trial, prior to any manipulations, ($2^{nd}$) individuals were weighted, ($3^{rd}$) control dominance trials between manipulations, and ($4^{th}$) dominance trial in which the top 20% ranked birds from the $1^{st}$ control from each loft had their iridescent plumage dampened resulting in decreased reflectance.

| Loft 1 | Steepness | Loft 2 | Steepness |
|---|---|---|---|
| $1^{st}$ Trial | 0.39 | $1^{st}$ Trial | 0.51 |
| $2^{nd}$ Trial | 0.46 | $2^{nd}$ Trial | 0.49 |
| $3^{rd}$ Trial | 0.37 | $3^{rd}$ Trial | 0.42 |

| | | | |
|---|---|---|---|
| $4^{th}$ Trial | 0.44 | $4^{th}$ Trial | 0.46 |

40
